# Quantifying microbial DNA in metagenomes improves microbial trait estimation

**DOI:** 10.1101/2024.06.20.599828

**Authors:** Raphael Eisenhofer, Antton Alberdi, Ben J. Woodcroft

**Author notes:** these authors contributed equally (Raphael Eisenhofer, Ben J. Woodcroft).

## Abstract

Shotgun metagenomics is a powerful tool for studying the genomic traits of microbial community members, such as genome size, gene content, etc. While such traits can be used to better understand the ecology and evolution of microbial communities, the accuracy of their estimations can be critically influenced by both known and unknown factors. One factor that can bias trait estimations is the proportion of eukaryotic DNA in a metagenome, as some bioinformatic tools assume that all DNA reads in a metagenome are non-eukaryotic. Here, we help resolve a recent debate about the influence of eukaryotic DNA in the estimation of average genome size from a global soil sample dataset using a new bioinformatic tool. Contrary to what was assumed, our reanalysis of this dataset revealed that soil samples can contain a substantial proportion of eukaryotic DNA (∼38.8%), which severely inflated average genome size estimates. We report that correcting for this bias significantly improves the statistical support for the negative relationship between average bacterial genome size and soil pH. These results highlight that metagenomes can contain large quantities of eukaryotic DNA, and that new methods that correct for this can improve microbial trait estimation.

## Main

Average genome size (AGS) is a trait that can be used to better understand microbial ecology and evolution[1]. For example, genome size is thought to reflect environmental and metabolic complexity, with smaller sizes associated with host dependency and reduced metabolic capacity[2,3]. In a 2023 article, Piton et al. observed a relationship between pH and AGS in soil samples, where larger average genome sizes were observed in lower pH samples[4]. Recently, Osmund et al. challenged this relationship, arguing that, since AGS was calculated as the ratio of the number of reads to the number of marker genes detected[5], AGS would be overestimated in soil samples containing eukaryotes. This could lead to systematic biases between different ecosystem types, with Osburn et al. arguing that the association between AGS and soil pH reported in the original study may be artefactual[6]. In response, Piton et al., while admitting this methodological limitation, argued that the alternative methodology proposed by Osburn et al. could be even more biased, and that soil does not contain substantial amounts of eukaryotic DNA[7]. Both research groups reached consensus on one point: “a perfect estimation of bacterial AGS using soil metagenomes is not yet possible”[7]. While not perfect, we argue that it is now possible to obtain fairly accurate estimations. We recently developed a tool that can account for such biases in AGS estimations[8], and we use it here to help settle this debate.

Our method, SingleM Microbial Fraction (“SMF”), can accurately estimate the fraction of bacterial and archaeal DNA from metagenomes[8]. Briefly, a marker-based approach is used to establish the coverage of each microbial taxon present. This coverage is multiplied by the genome size of that taxon, ultimately yielding an estimate of the community’s average genome size. This method was robust across a range of simulated datasets, and when applied to thousands of publicly available samples provided estimates consistent with other forms of validation. Critically, SMF’s average genome size estimates are not influenced by eukaryotic DNA—the major concern raised by Osburn et al. As part of our recent efforts to showcase the tool, we found that soil contains substantial quantities of eukaryotic DNA (median 31%, n = 4,160)[8]. This was in stark contrast to the assumption of an “extreme range of 4–9% eukaryotic base pairs” from Pitton et al.[7] (based on Supplementary Note 2 from[7]) — an assumption that was critical for their response to Osburn et al.

Leveraging SMF, we reappraised Piton et al.’s dataset to test whether eukaryote fractions varied across soil types, the extent of AGS overestimation due to this, and whether the original relationship between AGS and soil pH withstood correction. Substantial amounts of eukaryotic DNA were predicted (median 38.8%), representing a 9.7-to 4.3-fold underestimation by Piton et al.’s extreme range of eukaryotic base pairs[7] (**Figure 1A**). We also observed variation in eukaryotic DNA quantities both within and between soil environment types (**Figure 1A**). Our overall estimate of the mean AGS across all samples was 4.7 Mbp, 31% lower than originally estimated (6.8 Mbp). However, our mean AGS estimate was 36% higher than Osburn et al.’s (3 Mbp) estimate. This was likely because their approach only considered reads aligning to bacterial reference genomes, overlooking bacterial sequences not represented in databases[8,9]. We estimate that, as is common for soil metagenomes[10], only a small fraction of the microbial communities are represented by genomes at the species level (mean 0.5 ± 1.0% s.d.).

**Figure 1.**
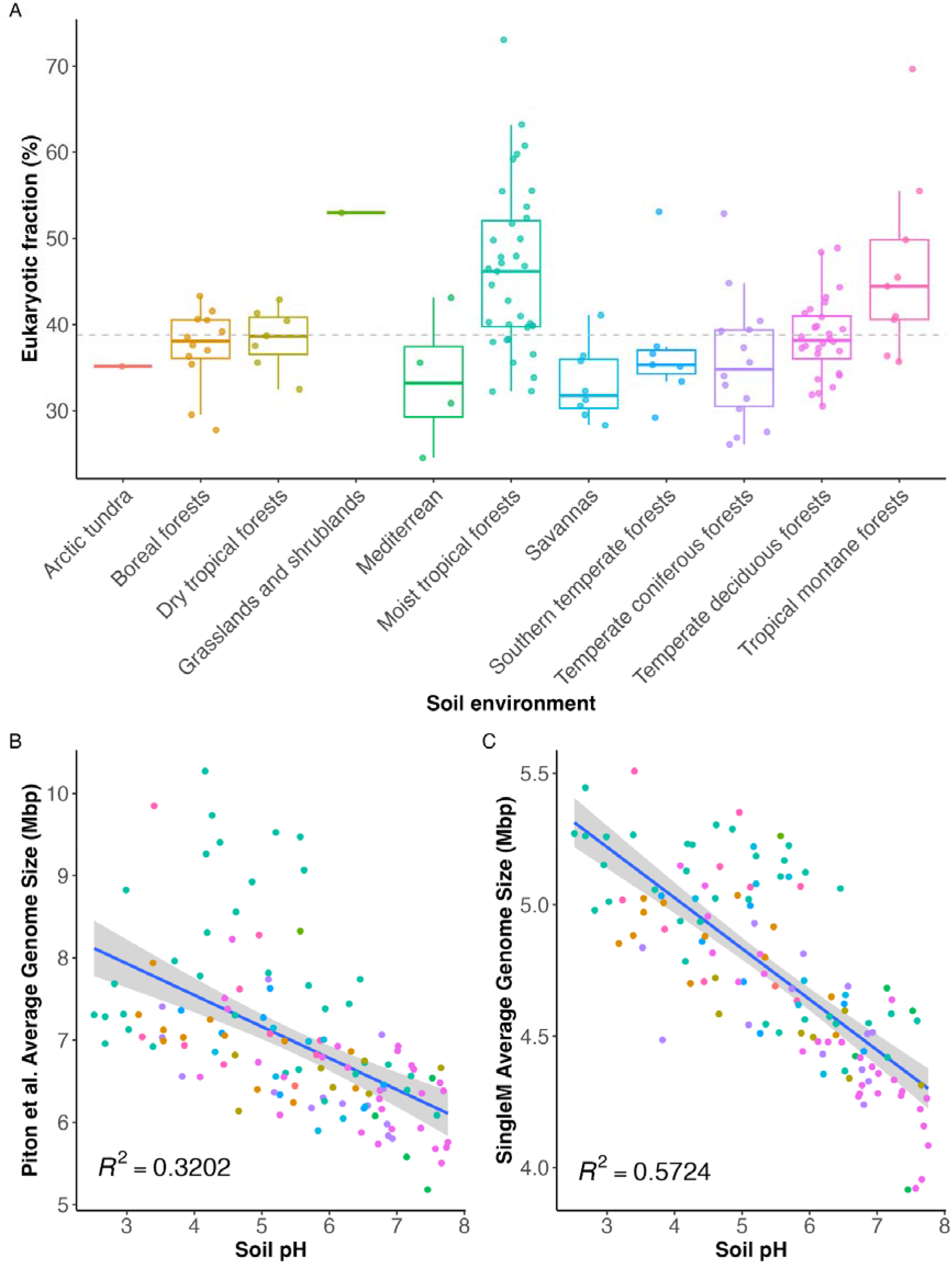
**A)** Variation in non-bacterial fraction between environment types. The percentage of eukaryotic base pairs (y-axis) across different soil environment types (x-axis. The dashed grey line represents the mean eukaryote fraction for the dataset. **B)** The relationship between Piton et al.’s AGS and soil pH. **C)** The relationship between SMF-corrected AGS and soil pH. *R* ^2^values and best-fit lines are from generalised linear models. *P* values < 0.0001.

Despite the variation of eukaryotic DNA observed in Piton et al.’s dataset and their overestimation of AGS, we found their original strong negative correlation between AGS and soil pH to be robust (**Figure 1B & 1C**). In fact, correcting for eukaryotic DNA abundance led to even stronger statistical support for this relationship (R^2^ 0.32 vs 0.57, **Figure 1C**). The relationship was also consistently observed within groups of soils of the same type (**SI Figure 1**). Finally, we also found the same correlation in an independent study of forest soils in China cited by Piton et al.[11], again with increased statistical support relative to the original study (R^2^ 0.42 vs 0.54, **SI Figure 2**).

The application of our newly developed tool helps to settle an ongoing scientific discussion that only a few months ago seemed unresolvable, and highlights the complexity and dynamism of the metagenomics research field. Our reappraisal of these data also highlights that eukaryotic DNA is currently a severely underestimated component of soil metagenomes. SMF can help identify and control for such biases in future bacterial (and archaeal) community trait estimations not only in soils, but other environments as well. While we have addressed one confounding factor, it is likely that metagenomic analyses still suffer from many unknown biases that will need to be identified and addressed in the near future. Only through critical scrutiny of our data and constructive discussions within the community can we ensure that metagenomics becomes the robust, broadly applicable tool we all aspire it to be.

## Methods

All data and code to reproduce the analyses and figures are available at the following GitHub repository: https://github.com/EisenRa/2024_soil_dark_matter_reply. To estimate the average genome sizes of samples from Piton et al.[4], datasets corresponding to each run accession were sourced from Sandpiper[10]. SingleM v0.17.0 “Archive OTU tables” of each sample were generated by merging those of the corresponding runs, and then re-assigned taxonomy using SingleM “renew” with a GTDB R214 reference metapackage (DOI: 10.5281/zenodo.11123537). The average genome size of each sample was then estimated using SingleM “microbial_fraction”. Soil pH values were obtained from https://doi.org/10.6084/m9.figshare.22620025. Data were imported, analysed and visualised in R[12] v4.3.2 using Tidyverse[13]. Generalised linear models were used to correlate average genome size to soil pH (glm() function, Gamma distribution).

To estimate the AGS in samples collected by Wang et al.[11], Reads from bioproject PRJNA986291 were downloaded using Kingfisher (https://github.com/wwood/kingfisher-download). Each run was then analysed using SingleM “pipe” and “microbial_fraction” as above. Measures of pH were those reported as the source data from Wang et al.’s Figure 2. *R*^2^ values were again calculated as above.

We note that reanalysis was undertaken considering bacteria to be the primary constituents of the microbial community, since Archaea were found to be in very low abundance (mean 1.2 ± 0.7% s.d. according to SingleM). Specifically, no attempt was made to separate out the bacterial and archaeal components of the community, and all genome size calculations reported here are those of the total (bacterial and archaeal) microbial community.

## Acknowledgements

B.J.W. was supported by Australian Research Council Future Fellow (#FT210100521), Discovery Project (#DP230101171) and the EMERGE Biology Integration Institute, funded by the NSF Biology Integration Institutes Program, Award no. 2022070 grants. A.A. acknowledges the Danish National Research Foundation award DNRF143 ‘A Center for Evolutionary Hologenomics’ and the Carlsberg Foundation grant CF20-0460.

## Supplementary information

**Supplementary Figure 1.**
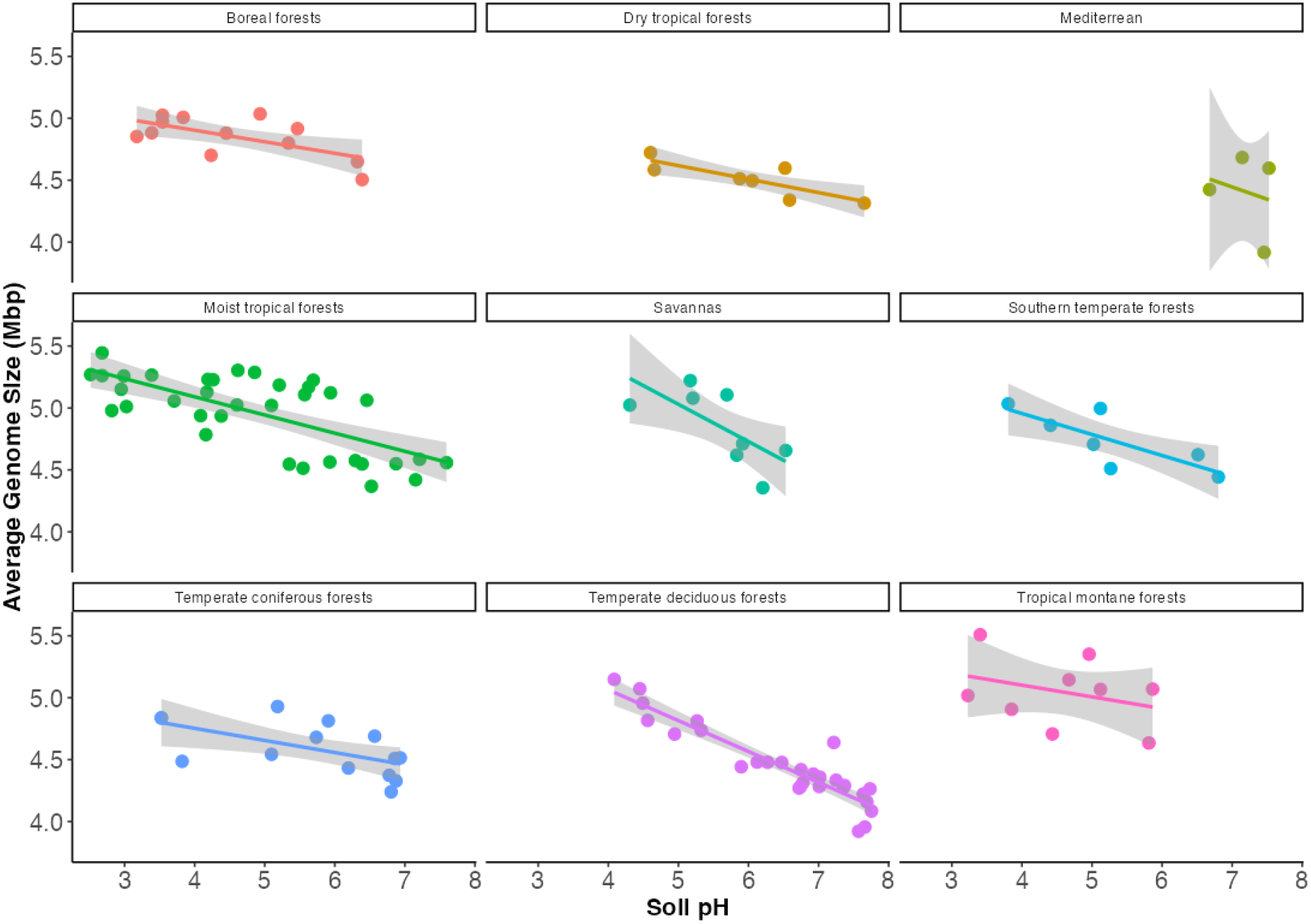
The relationship between corrected AGS and soil pH faceted by soil type. Best-fit lines are from generalised linear models.

**Supplementary Figure 2.**
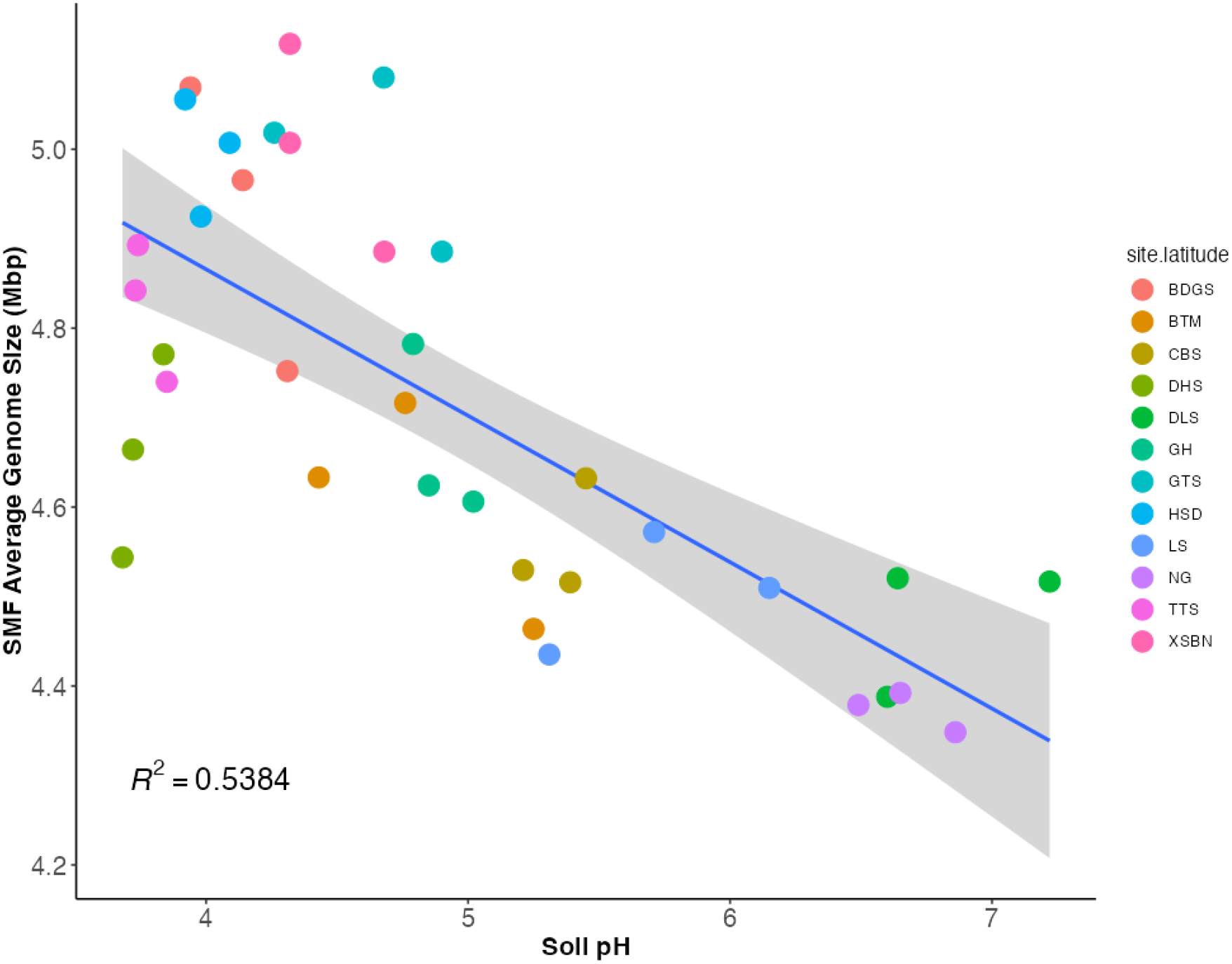
Reanalysis of Wang et al. 2023 soil samples. The relationship between corrected AGS (y-axis) and soil pH (x-axis). *R*^2^values and best-fit lines are from generalised linear models. *P* < 0.0001.

